# Integrative conformational ensembles of Sic1 using different initial pools and optimization methods

**DOI:** 10.1101/2022.04.01.486785

**Authors:** Gregory-Neal W. Gomes, Ashley Namini, Claudiu C. Gradinaru

**Affiliations:** Department of Physics, University of Toronto, Toronto, Ontario, M5S 1A7, Canada; Department of Chemical & Physical Sciences, University of Toronto Mississauga, Mississauga, Ontario, L5L 1C6, Canada

**Keywords:** IDPs, smFRET, NMR, SAXS, ENSEMBLE, BME, contact maps, hydrogen bonds, pi interactions

## Abstract

Intrinsically disordered proteins play key roles in regulatory protein interactions, but their detailed structural characterization remains challenging. Here we calculate and compare conformational ensembles for the disordered protein Sic1 from yeast, starting from initial ensembles that were generated either by statistical sampling of the conformational landscape, or by molecular dynamics simulations. Two popular, yet contrasting optimization methods were used, ENSEMBLE and Bayesian Maximum Entropy, to achieve agreement with experimental data from nuclear magnetic resonance, small-angle X-ray scattering and single-molecule Förster resonance energy transfer. The comparative analysis of the optimized ensembles, including secondary structure propensity, inter-residue contact maps, and the distributions of hydrogen bond and pi interactions, revealed the importance of the physics-based generation of initial ensembles. The analysis also provides insights into designing new experiments that can maximally discriminate among the optimized ensembles. Overall, differences between ensembles optimized from different priors were greater than when using the same prior with different optimization methods. Generating increasingly accurate, reliable and experimentally validated ensembles for disordered proteins is an important step towards a mechanistic understanding of their biological function and involvement in various diseases.

## INTRODUCTION

Important biological functions performed by intrinsically disordered proteins (IDPs), such as cell signaling and regulation (Dyson and Wright, 2005; Forman-Kay and Mittag, 2013; Oldfield and Dunker, 2014), are mediated by their interesting and nonrandom structural properties. Conversely, their dysfunction or pathological aggregation is accompanied or preceded by aberrations in these structural properties (Uversky, 2015). Describing the molecular features of IDPs at atomistic resolution would therefore provide valuable mechanistic insight into how IDPs (mal)function. Molecular dynamics (MD) simulations have recently attempted to fill this gap, including development of new force fields to accurately model disordered proteins (Best et al., 2014; Rauscher et al., 2015). However, a unique parametrization of force fields suitable for modelling IDPs is yet to emerge, and atomistic-level simulations over biologically relevant timescales remain computationally expensive. Alternatively, disordered proteins can be represented by a conformational ensemble, which is a finite set of 3D structures with corresponding statistical weights. These ensembles are commonly determined by re-weighting or selecting a subset from an initial pool of conformations according to a protocol which optimizes agreement with various experimental data, while considering experimental uncertainties and avoiding overfitting (Bonomi et al., 2017; Bottaro et al., 2020; Jensen et al., 2014; Köfinger et al., 2019; Krzeminski et al., 2013; Lazar et al., 2021; Leung et al., 2016; Lincoff et al., 2020; Orioli et al., 2020).

Recent and rapid progress in the field of protein disorder necessitates a re-examination of the ensemble determination process. Mutual consistency and complementarity have been demonstrated for the three most commonly used structural techniques for IDPs: Small Angle X-Ray Scattering (SAXS), Nuclear Magnetic Resonance (NMR) and single-molecule Förster Resonance Energy Transfer (smFRET) (Aznauryan et al., 2016; Delaforge et al., 2015; Fuertes et al., 2017; Gomes et al., 2020; Lincoff et al., 2020; Naudi-Fabra et al., 2021; Voithenberg et al., 2016). Technological advances and efforts to standardize data collection and reporting have also been made for SAXS (Martin et al., 2020), smFRET (Hellenkamp et al., 2018; Lerner et al., 2021) and NMR (Alderson and Kay, 2021; Dyson and Wright, 2021, 2019). Improvements in the accuracy of MD force fields, which correct earlier bias toward overly compact IDP conformations (Best et al., 2014; Huang et al., 2017; Rauscher et al., 2015; Robustelli et al., 2018), have advanced their use for generating initial pools of conformers. Protocols for calculating ensembles (Bottaro et al., 2020; Köfinger et al., 2019; Leung et al., 2016; Lincoff et al., 2020) and for predicting experimental data from structures (Crehuet et al., 2019; Dimura et al., 2020; Henriques et al., 2018; Kalinin et al., 2012; Pesce and Lindorff-Larsen, 2021; Tesei et al., 2021) continue to be developed and refined. As a result of all these developments, the repository of IDP ensembles validated by agreement with experimental data, the Protein Ensemble Database, has recently undergone a major update (PED 4.0) (Lazar et al., 2021).

The high conformational entropy and extreme conformational dynamics of IDPs, however, remain the major challenges to this overall project. Experimental data provide time- and ensemble-averaged structural information which is affected by random and possibly systematic errors. As such, the number of degrees of freedom necessary to specify an ensemble of atomic resolution structures is inherently much larger than the number of experimentally determined structural restraints. Ensemble calculation is therefore a mathematically ‘ill-posed’ or ‘underdetermined’ problem that always has more than one solution (Bonomi et al., 2017; Bottaro et al., 2020; Lazar et al., 2021; Marsh and Forman-Kay, 2012).

Differences in how ensembles are determined, such as how an initial ensemble is generated and which ensemble optimization algorithm is used, lead to further proliferation in the number of possible solutions for the same experimental dataset. Trivially, these ensembles are distinct as they are composed of different protein conformations. However, whether these differences are significant or not remains unclear, and it will require a quantitative comparison of their impact on inferences about sequence-structure or structure-function relationships. Understanding this variability in calculated ensembles for the same system is particularly important given the renewed efforts of PED 4.0 to curate high quality ensemble structural data (Lazar et al., 2021).

To probe the intrinsic variability of this under-determined process and evaluate its effect on sequence-structure-function relationships, we examined ensembles generated from different conformational priors and using different modelling methodologies. Broadly, prior ensembles can be generated using either: *(i)* MD simulations, which use physics-based force fields to generate Boltzmann-weighted ensembles; or *(ii)* statistical coil approaches, which use extensive (un)biased sampling of the complete conformational phase space. Here, we selected two MD priors, Amber ff03ws (Best et al., 2014) (a03ws) and Amber 99SBdisp (Robustelli et al., 2018) (a99SBdisp), and a statistical coil prior generated by TraDES (Feldman and Hogue, 2002), TraDES-SC.

A03ws is a force field in which the protein-water interactions in the a03w protein forcefield were rescaled by a constant factor to produce more realistic dimensions of denatured and intrinsically disordered proteins (Best et al., 2014). A99SBdisp is a recently developed force-field intended to provide accurate descriptions of both folded and disordered proteins (Robustelli et al., 2018). In a recent benchmarking study, a03ws was shown to produce global dimensions agreeing with experiment, but at the expense of residual secondary structure propensity of IDPs or stability of folded proteins (Robustelli et al., 2018). In the same study, a99SBdisp accurately described both ordered and disordered states, including global dimensions of many IDPs. However, for larger IDPs with more hydrophobic sequences (*α*-synuclein, *N*_*TAIL*_, Sic1), a99SBdisp showed a bias toward overly compact global dimensions. In contrast, TraDES generates all-atom conformations in which the only physics-based interactions are excluded-volume (Feldman and Hogue, 2002).

We have selected two popular, but contrasting modeling methodologies: the Bayesian Maximum Entropy (BME) (Bottaro et al., 2020) approach and ENSEMBLE (Krzeminski et al., 2013). Although there are many specific differences between these methodologies, the major distinction is in the treatment of the prior ensemble and of experimental and prediction errors. The BME approach produces the minimum perturbation to the prior ensemble (i.e., maximum relative Shannon entropy with respect to the prior) such that it fits the experimental data, with experimental and prediction errors accounted for in a Bayesian framework. The ENSEMBLE approach, in contrast, places no restriction on the deviation from the prior ensemble while minimizing pseudo-energy terms quantifying disagreement with experimental data. These pseudo-energy terms are typically harmonic potentials with preset scaling and target energies.

We focus here on the N-terminal 90 residues of the full-length disordered protein Sic1 (henceforth Sic1) which has been extensively characterized by NMR, SAXS and smFRET experiments (Gomes et al., 2020; Liu et al., 2014; Mittag et al., 2008, 2010) and for which we have recently determined ensembles using the ENSEMBLE method (Gomes et al., 2020). In their benchmarking study and to test their recently developed a99SBdisp forcefield, Robustelli et al., produced long-timescale (30 *μ*s) simulations of Sic1 using a03ws and a99SBdisp (Robustelli et al., 2018). The authors have kindly provided these simulations to be used as prior ensembles in our calculations. Importantly, Sic1 was in the test set and not in the training set for developing a99SBdisp. The extensive experimental characterization and molecular modelling of Sic1 make it an ideal case for benchmarking both future force-field developments and ensemble modelling.

To compare the six resulting Sic1 ensembles (two modelling methodologies, three prior ensembles), we compare their secondary structure propensities and the mean inter-residue distances in the ensembles (tertiary structural propensities). The three prior ensembles differ in how physics-based interactions are considered and the two modelling methodologies differ in how information is retained from the prior ensemble. Accordingly, we also compare the nature of the molecular interactions (hydrogen bond and pi-pi) that may be present in the calculated ensembles.

## RESULTS and DISCUSSION

### Bayesian Maximum Entropy

The BME method is equivalent to minimizing the objective function 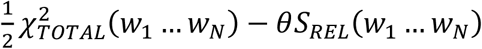 where the *w*_*i*_ are the optimized weights for each conformer in the prior ensemble. Here, 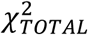 quantifies the total agreement with all experimental data points and 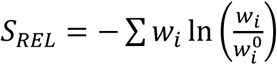 is the relative entropy which quantifies the deviation from the initial weights, 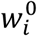, in our case all equal to 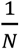. The hyperparameter *θ* balances the confidence in the prior with respect to that of the experimental data and it is determined by tuning (discussed below), given that the combined uncertainty in the experimental data, the calculated data, and the prior ensemble is not known accurately. For more details on the theory behind BME, the reader is referred to the original author’s publications (Bottaro et al., 2020; Orioli et al., 2020) and equivalent or similar approaches (Köfinger et al., 2019; Leung et al., 2016).

In this work, we use SAXS, chemical shifts (CS) and smFRET data (between residues −1C and 90C, probing approximately the end-to-end distance) as restraints, and so 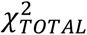 is the sum of the individual non-reduced *χ*^2^s. In principle, the relative weights of each experiment could be determined accurately if the following could be determined accurately: *(i)* the degrees of freedom for each experiment; *(ii)* the statistical *and* systematic experimental uncertainties; and *(iii)* the statistical *and* systematic uncertainties in the “forward calculation” (calculation of experimental observables from structures). In this case, 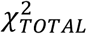 would simply be the sum of each experiment’s individual non-reduced *χ*^2^, i.e., 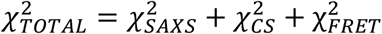. However, in practice *(i)*—*(iii)* are not possible to determine accurately, as discussed below. In the absence of a corrective, datatypes with many datapoints (i.e., SAXS and CSs) would overwhelm those with one or a few observations (i.e., smFRET). To compensate for the undue influence of SAXS and CS relative to FRET on the 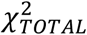 we introduced and determined a weighting factor Ω such that 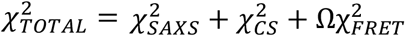 (SI section 1, Fig. S1, S2). Briefly, increasing Ω from Ω = 1 to Ω ≈ 75 improves the fit to the smFRET data and a set of independent validation data (see below), without worsening the SAXS fit, and with only marginally more reweighting.

Figure 1A-C shows how *θ* was determined for the three prior ensembles: a03ws, a99SBdisp, and TraDES-SC, respectively. Lower values of *θ* result in greater agreement with experiment (lower 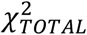) but with greater deviation from the prior ensemble, which is quantified by a lower effective number of conformations used in the posterior ensemble, *N*_*eff*_ = exp(*S*_*REL*_). Although better agreement with experiments could be achieved by letting *θ* → 0, within the BME framework this would *(i)* ignore uncertainties in the experimental and calculated values and *(ii)* disregard information about molecular interactions encoded in the priors, e.g., the physics of the force fields in the MD priors. Enforcing too tight an agreement with the set of restraining data can also lead to overfitting. To verify if overfitting occurs and find the optimal value of *θ*, we assessed the agreement of the ensembles with six PRE experiments (*m*_*PRE*_ ≈ 400 data points), which were not used as input for reweighting. As such, we scanned through different values of *θ* and simultaneously monitored 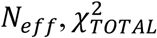 and the PRE score calculated using DEER-PREdict (Tesei et al., 2021).

**Figure 1:**
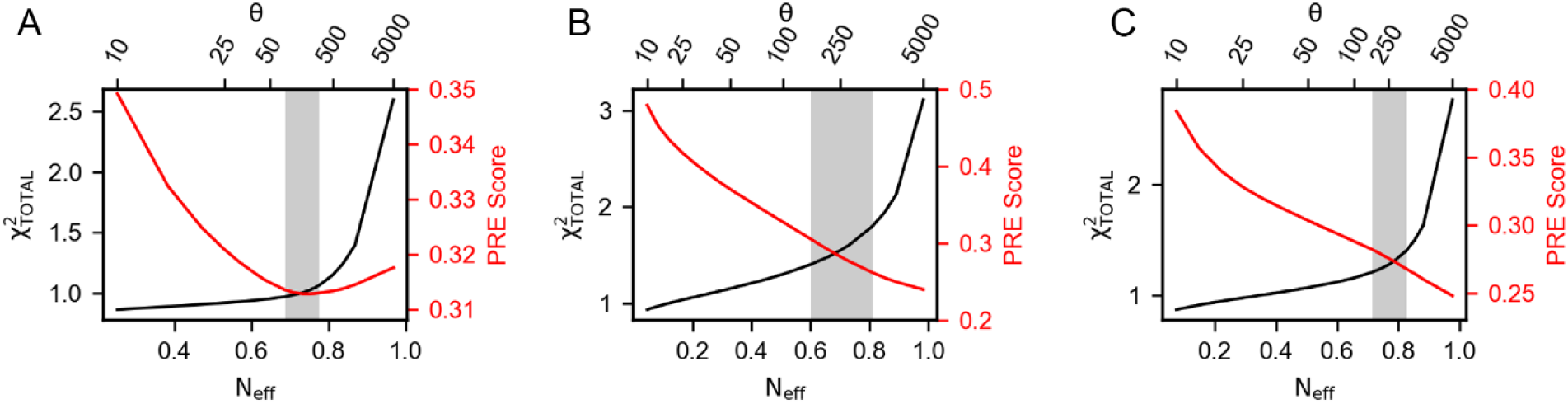
Optimization of Sic1 conformational ensembles using BME and different initial pools: a03ws (**A**) and a99SBdisp (**B**), and TraDES-SC (**C**). For each value of *θ* (top x-axis), agreement with the restraining data (SAXS, CS and smFRET) is measured by 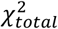 (black). Agreement of the posterior ensemble with PRE data (PRE Score, red) serves as validation. For a03ws (**A**), *θ* was chosen to minimize disagreement with the validation data; for a99SBdisp (**B**) and TraDES-SC (**C**), *θ* was chosen at the “elbow” in the 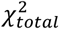 identified by L-curve analysis. In each case, a range of acceptable *θ* was evaluated near the optimal values (grey shaded region, Table 1).

For all three priors, there is an initial region in which lower values of *θ* substantially decrease 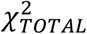 with only small decreases in *N*_*eff*_, followed by a region in which small decreases in 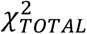 are accompanied by substantial decreases in *N*_*eff*_. We therefore use L-curve analysis (Hansen and O’Leary, 1993; Orioli et al., 2020) to identify a useful region of *θ* corresponding to the “elbow” regions in the *N*_*eff*_ vs 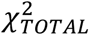 plots. For a03ws, the agreement with PRE data initially improves (possibly because of distances shared by the smFRET restraint and PRE validation), then it worsens as *θ* is set beyond the elbow region. In contrast, for a99SBdisp and TraDES-SC priors, agreement with the PRE validation data monotonically worsens as *θ* decreases. We hypothesize that this is a consequence of enforcing the SAXS restraint on relatively compact prior ensembles, as discussed below.

**Table 1.**
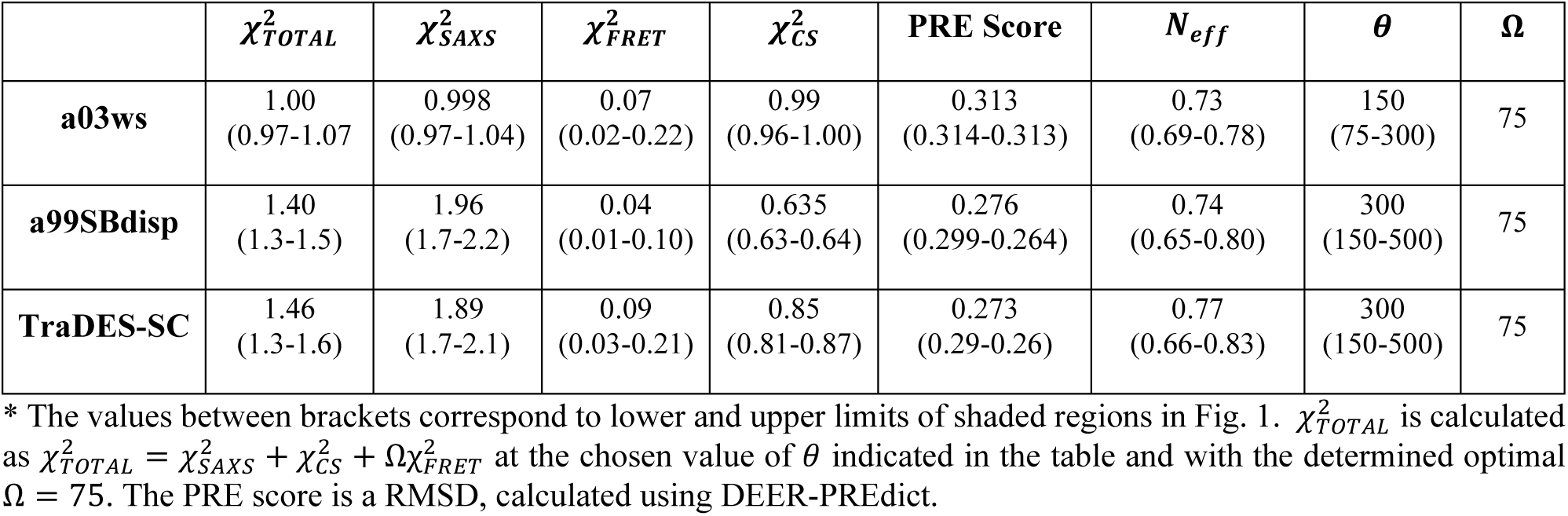
Optimization parameters of BME-calculated Sic1 ensembles*

Due to the *r*^−6^ averaging of nuclear-electron distances in PRE measurements, the ensemble averages are dominated by contributions from compact conformers (Ganguly and Chen, 2009). As such, the presence of few compact conformers can satisfy the PRE data (Ganguly and Chen, 2009), and decreasing the weight of these conformers to satisfy the SAXS restraint worsens agreement with the PRE validation. In contrast to a03ws, which is already in good agreement with the SAXS data before re-weighting, a99SBdisp and TraDES-SC are more compact, with fewer conformations that are expanded above the experimental radius of gyration, *R*_*g*_ (SI, Fig. S4). As a result, deriving ensembles for a99SBdisp and TraDES-SC that agree with the SAXS data^46^ involve significant re-weighting of the prior ensembles by reducing the weight of compact conformations and increasing the weight of expanded conformations.

For further analysis, we selected *θ* resulting in the minimum of the PRE validation score for a03ws, and *θ* corresponding to the “elbow” region of the 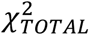 vs. *N*_*eff*_ plots for a99SBdisp and TraDES-SC (Fig. 1). Table 1 shows the reduced 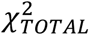, the individual restraining data reduced *χ*^2^s, the PRE validation score, and the *N*_*eff*_ for each prior at their corresponding optimal *θ* values. Although re-weighting improved the agreement of a99SBdisp and TraDES-SC ensembles with the SAXS data, it is not possible to improve it further without substantially deviating from the prior ensemble (very low *N*_*eff*_) and incurring overfitting (e.g., poor PRE validation performance). All three posterior ensembles agree well with the smFRET data; for a03ws, this is a result of re-weighting, while prior a99SBdisp and TraDES-SC ensembles were already in reasonably good agreement. However, the end-to-end distance measured by smFRET has a high restraining strength, as in its absence the global expansion dictated by the SAXS data would lead to anomalously expanded ensembles for a99SBdisp and TraDES-SC (Fuertes et al., 2017; Gomes et al., 2020).

### ENSEMBLE

The ENSEMBLE method (Krzeminski et al., 2013) minimizes a total pseudo-energy, which is the weighted sum of each individual experiment’s pseudo-energy, wherein lower energies correspond to better agreement with experimental restraints. To perform this minimization, ENSEMBLE employs a switching Monte-Carlo algorithm within a simulated annealing protocol to select subsets of conformers from the initial ensemble. The optimization terminates when all experimental restraints are below their respective target energies that are set by default in ENSEMBLE (Krzeminski et al., 2013). The relative weights of different experiments are adjusted during optimization, with increased weight given to experiments that are above their target energies. We perform five independent ENSEMBLE calculations with 100 conformations and combine the results to form ensembles with 500 conformations, based on previous calculations (Gomes et al., 2020; Marsh and Forman-Kay, 2012). This ensemble size balances between the concerns of overfitting and underfitting and ideally, structural features resulting from overfitting should be averaged out in independent calculations. When applying ENSEMBLE to Sic1, we used SAXS, CS, and PRE data as experimental restraints, and reserved the smFRET data as a validation. Allocating the experimental data into restraints and validation identically for both optimization methods is not currently possible since ENSEMBLE and BME accommodate different experimental data types.

Figure 2 shows typical ENSEMBLE pseudo-energy minimizations for all three priors as a function of the number of Monte-Carlo trials. Note that because the ENSEMBLE optimization is stochastic, no two trajectories will be identical. Each pseudo-energy is normalized by its ENSEMBLE-defined target energy, such that a value less than one is considered “fit” by the program (gray shaded region). The smFRET validation is shown as a solid blue line with the right-hand axis, with the blue-shaded region corresponding to the experimental FRET efficiency ⟨*E*⟩_*exp*_ and its uncertainty. Ensembles with a FRET value within the blue region are within 1*σ* of the experimental value.

**Figure 2:**
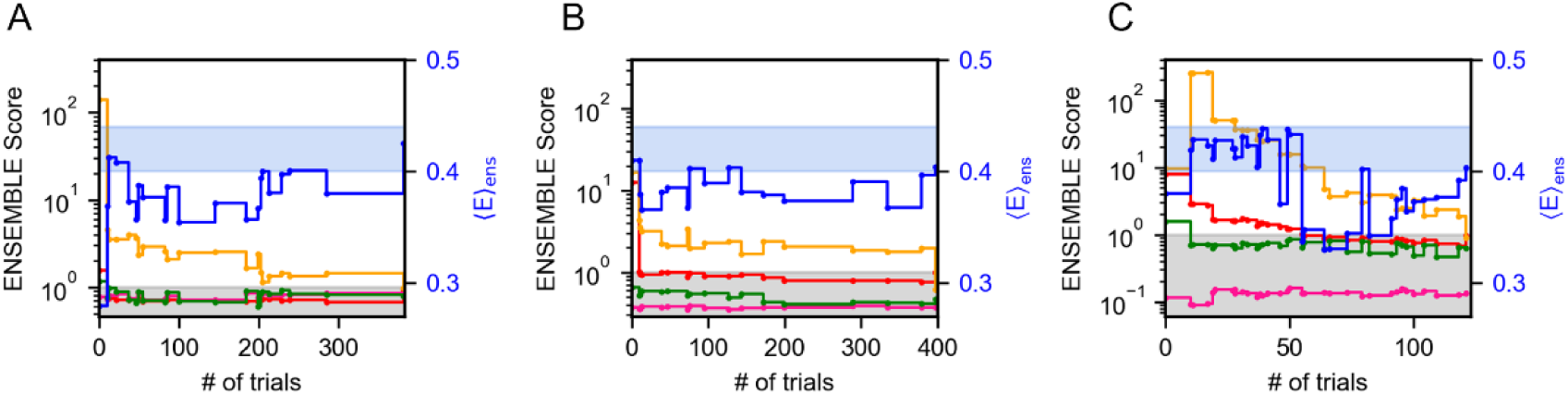
Optimization of Sic1 conformational ensembles using ENSEMBLE and different initial pools: a03ws (**A**) and a99SBdisp (**B**), and TraDES-SC (**C**). Individual restraint pseudo-energies are normalized by their ENSEMBLE-defined target energy, such that a value less than one is considered “satisfied” (gray shaded region). Shown here are typical trajectories from the ENSEMBLE optimization of each prior (a03ws, a99SBdisp, TraDES-SC) using the following restraints: SAXS (red), chemical shifts (alpha - green, beta - magenta), and PRE (yellow). smFRET is used as an external validation, with the blue-shaded region showing the measured efficiency, ⟨*E*⟩_exp_, and its uncertainty.

For a03ws (Fig. 2A), energy minimization is largely focused on improving the agreement with the PRE data, whereas the trial ensembles agree with the CS and SAXS data either initially or after relatively few trials. For a99SBdisp (Fig. 2B), the initial disagreement with the PRE data is less than for a03ws, though the initial disagreement with the SAXS data is greater. However, in relatively few trials the SAXS data is fit, and further energy minimization is focused on the PRE data. In contrast to a03ws and a99SBdisp, which are new MD force fields designed to accurately describe IDPs, TraDES-SC (Fig. 2C) only accounts for excluded volume and random propensities for varying secondary structure (hence, statistical coil). Unsurprisingly, the TraDES-SC ensemble initially disagrees with most of the experimental data. Optimization first reduces the SAXS restraint below its target energy, before finally fitting the PRE data.

For all ENSEMBLE calculations, the PRE restraint was the last to be fit below its target energy, while the CS data was fit either initially or within the first few trials. This suggests that CSs are a comparatively easy experimental restraint to meet, perhaps because of the comparatively large CS calculator uncertainties. Consequently, the secondary structure propensities of the optimized ensembles will be largely dictated by the propensities of the prior ensembles (see below). As shown in Fig. 2, trial ensembles which fit the SAXS data but not the PRE data have overly expanded end-to-end distances resulting in ⟨*E*⟩ < ⟨*E*⟩_*exp*_. Jointly fitting the SAXS and PRE data places strong restraints on the end-to-end distance distribution, and consequently on ⟨*E*⟩. This reinforces the conclusions drawn by Gomes et al., which were made using only the TraDES-SC prior ensemble, and emphasizes the inability of SAXS to determine specific inter-residue distances, except in the case of ideal polymer models (Gomes et al., 2020).

Table 2 shows the mean and standard deviation of the non-normalized ENSEMBLE total energy upon termination for the five independent trials. Although ENSEMBLE minimizes an ENSEMBLE-defined energy term for each experimental data type, Table 2 shows the reduced *χ*^2^ for SAXS and CS data (C*α* and C*β* combined) to facilitate comparison of the fits with those done by BME. ENSEMBLE optimization considers PRE restraints as 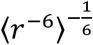 distance restraints and approximates the electron location of the paramagnetic probe to the position of the C*β* atom of the spin-labelled residue (Krzeminski et al., 2013). In Table 2, the PRE score is the RMSD between the calculated and experimental PRE intensity ratios using the more accurate rotamer library approach, DEER-PREdict (Tesei et al., 2021), which was also used to calculate BME validation scores (see above). Additionally, the agreement with the smFRET validation data is reported using a z-test.

**Table 2.** Optimization parameters of ENSEMBLE-calculated Sic1 ensembles

| | **ENSEMBLE**<br>**energy** | $\chi^2_{SAXS}$ | **PRE score** | $\chi^2_{CS}$ | **z-test**<br>**FRET\*** | **Number of trials** |
| --- | --- | --- | --- | --- | --- | --- |
| **a03ws** | $335 \pm 17$ | 0.985 | 0.285 | 0.782 | 0.73 | $327 \pm 102$ |
| **a99SB-disp** | $302 \pm 6$ | 0.976 | 0.264 | 0.478 | 0.67 | $452 \pm 261$ |
| **TraDES-SC** | $272 \pm 15$ | 0.986 | 0.201 | 0.324 | 1.03 | $141 \pm 33$ |

\* $z_{\langle E \rangle} = |\langle E \rangle_{ens} - \langle E \rangle_{exp}| / \sigma$ . Uncertainty in ENSEMBLE energy and Number of trials is calculated as the standard deviation of the five independent ENSEMBLE optimizations. $\chi^2_{SAXS}$ , PRE score and $\chi^2_{CS}$ are calculated for the composite 500 conformer optimized ensemble. PRE scores for the final ensembles are calculated using DEER-PREdict to calculate an RMSD, as for the BME validation scores; however, during optimization the ENSEMBLE’s native PRE module was used.

Interestingly, the ENSEMBLE-optimized TraDES-SC ensemble is in better agreement with the PRE and CS data than the ENSEMBLE-optimized a03ws and a99SBdisp ensembles. This may be due to the much larger conformational diversity in the TraDES-SC initial pool. When optimizing for a03ws and a99SBdisp, no new conformations are generated, and ENSEMBLE must select from the fixed initial pool of MD-generated conformers. For TraDES-SC, we used ENSEMBLE’s built-in conformer generation and management (Krzeminski et al., 2013), in which new conformations are regularly replenished using TraDES. The conformer management algorithm favors conformers that have been selected fewer times in Monte-Carlo trials. Moreover, conformations in the MD prior ensembles will naturally have some degree of structural correlation as they are generated by the system’s time-evolution. The increased sampling of conformational space for the ENSEMBLE optimized TraDES-SC ensemble might explain the more rapid approach to the final solution (fewer trials, see Table 2), and the lower PRE score and 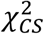 when compared to the optimization using MD priors.

### Secondary structure propensity (SSP)

Secondary structures of proteins are defined by specific patterns of hydrogen bonds, dihedral angles and other geometrical restraints. Based on the continuously expanding library of 3D structures in the Protein Data Bank (PDB, www.rcsb.org), various algorithms were developed to classify and predict secondary structure motifs in proteins (Reeb and Rost, 2019). Define Secondary Structure of Proteins (*DSSP*) annotates secondary structure elements to one of eight possible states and groups them into three classes: *helical* (α-, 3_10_- and π-helices), *strand/extended* (β-bridges and β-bulges) and *loop/coil* (turn, bend and other) (Kabsch and Sander, 1983; Touw et al., 2015). Figure 3 shows the DSSP distributions of the three classes of secondary structure (helical, extended and coil) for 6 optimized Sic1 ensembles (2 methods and 3 priors).

**Figure 3.**
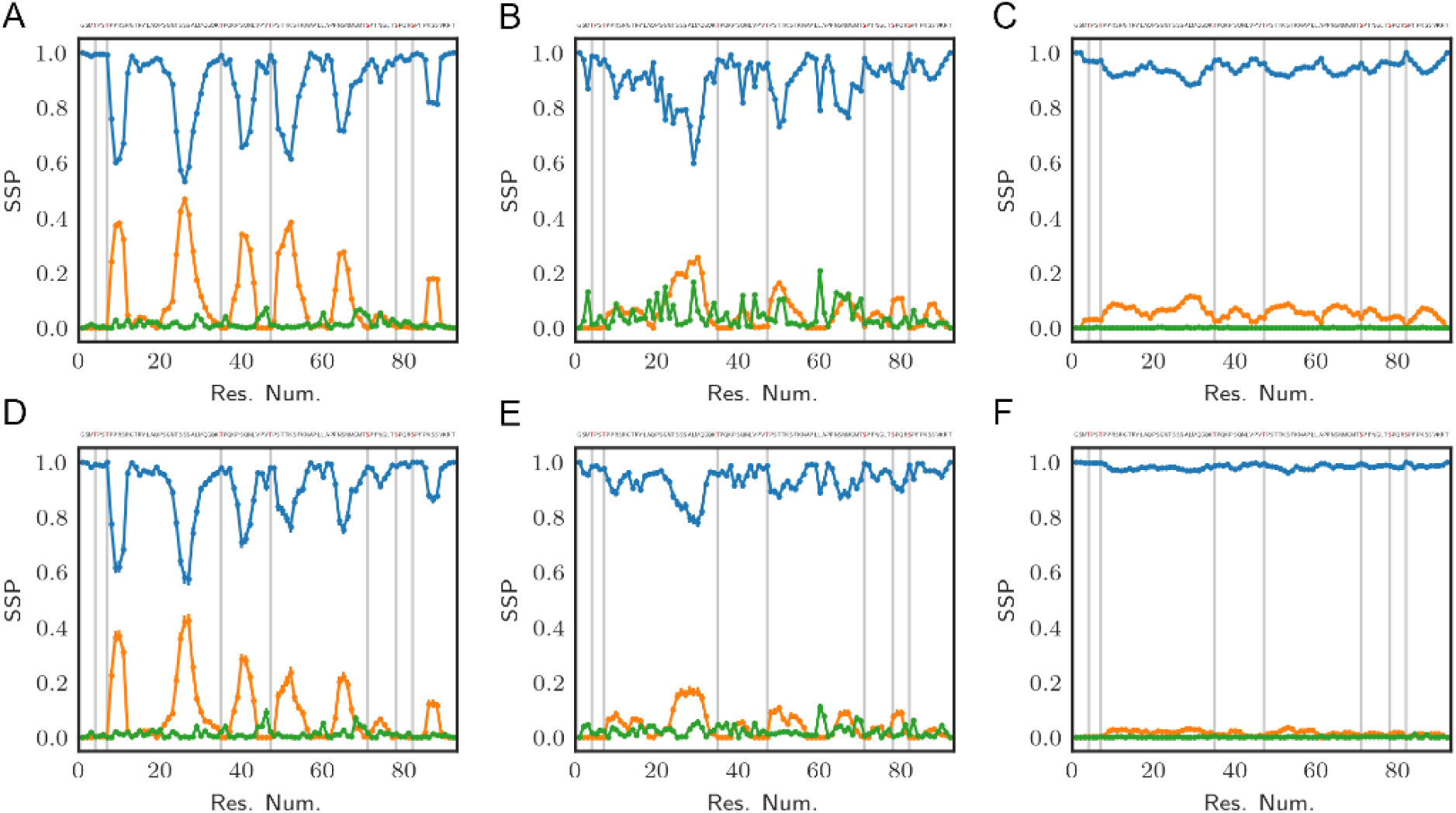
Secondary structure propensity (SSP) of optimal Sic1 ensembles using the DSSP algorithm (Kabsch and Sander, 1983; McGibbon et al., 2015). The ensembles were calculated using BME (A-C) or ENSEMBLE (D-F), and different initial ensembles: a03ws (A, D) a99SBdisp (B, E), and TraDES-SC (C, F). For each ensemble, secondary structure elements shown are *coil* (blue), *alpha-helix* (orange), and *beta/extended* (green). Error bars were calculated using bootstrapping. Phosphorylation sites (S and T residues) are shown in the Sic 1 sequence at the top of each panel in red, and as grey vertical lines in each panel.

The TraDES-SC ensembles stand out as almost exclusively consisting of coil structures (>90% for BME, >95% for ENSEMBLE), with essentially null fraction of extended elements, and at most 10% of helical fraction quasi-uniformly distributed throughout the sequence (Fig. 3C, F). At the other end of the spectrum, the a03ws ensembles exhibit much larger helical propensities at the expense of the coil fraction. There are six 5-10 residue helical patches distributed throughout the sequence around serine residues, with propensities ranging from ~0.2 near the C-terminus (S87) to ~0.5 around S26 (Fig. 3A, D). The ENSEMBLE optimization allows the experimental restraints to act on the prior more aggressively, leading to a reduction of the helical propensities by ~0.05 for each patch, although the sequence distribution is preserved. While the extended structure propensities are higher than when using the TraDES-SC prior, they do not exceed 0.05 and appear as short patches interleaved with the larger helical patches.

The ensembles calculated from the a99SBdisp prior reveal an intermediate picture between the two other cases (Fig. 3 B, E). The 6 helical patches present in the a03ws ensembles are still present here, albeit at a reduced propensity (~0.1 to ~0.25), with the BME method again exhibiting slightly larger values. Notably, a higher beta/extended propensity is observed at various points throughout the sequence, with the BME ensemble showing more of them and with larger values (~0.2) than the one obtained by ENSEMBLE (~0.1).

To a large extent, the differences in the DSSP maps reflect inherent differences in the structural ensembles used as priors. The impact of the optimization method on secondary is limited, with BME (by design) effecting a smaller bias of the prior than ENSEMBLE. TraDES-SC, which we used in a recent study of Sic1 (Gomes et al., 2020), is the least sophisticated prior of the three studied here, as it includes only excluded-volume interactions between chain residues. It is not surprising that imposing averaged size and chemical shift restraints on this ensemble cannot create “de novo” secondary structure. The chance of bringing patches of residues within hydrogen bond contact with peptide backbone forming specific dihedral angles is infinitesimally small, especially given the level of imprecision in the back calculators and the error margins of the experimental values.

Robustelli et al benchmarked several MD force fields to describe the properties (size, secondary structure, etc.,) of both folded and disordered proteins, including Sic1 (Robustelli et al., 2018). Among those, a03ws, which empirically optimized protein-water dispersion interactions for disordered protein (Best et al., 2014), reproduced the *R*_*G*_ most accurately and exhibited relatively large helical propensities. However, in addition to experimentally observed helices, it also populated regions where helical propensity were not observed experimentally. As such, it is not surprising that the a03ws prior contains the highest fraction of helicity of all the priors used here. On the other hand, a99SBdisp is a force field with modified Lennard-Jones parameters (Nerenberg et al., 2012) and optimized torsion angles and van der Waals parameters, which achieved best scores in matching the experimental observables for both folded and unfolded proteins in the benchmark set (Robustelli et al., 2018). For disordered proteins (e.g., α-synuclein) a99SBdisp shows less helical propensity than a03ws, a trend that is also observed for Sic1. As mentioned above for TraDES, the impact of experimental restraints on biasing the secondary structure in ensemble calculations is minor (a slight decrease for a03ws and a slight increase for a99SBdisp). The physics model of the prior, i.e., the force field parametrization, is by far the most important factor that drives formation of stable/transient secondary structure motifs.

In the case of Sic1, it is worth comparing the DSSP maps of the optimized ensembles with the SSP scores calculated using chemical shift data^59^. Three of the six helical patches observed in the DSSP maps are also present in the SSP map (around res. #26, 50 and 65), however, in contrast to SSP, regions of extended secondary structure were not significantly populated by DSSP for any of the 6 cases examined. Notably, each of the seven phosphorylation sites (indicated by vertical grey bars in Fig. 3) lies outside the helical patches, in the coil regions of Sic1. This may ensure access of kinase enzymes to these sites, and favor a binding model in which the multiple CPD sites in Sic1 engage the single receptor site of Cdc4 in a fast dynamic equilibrium. On the other hand, the ubiquitination sites in Sic1 (Lys 32, 36, 50, 53, 84 and 88) must bridge a 64 Å between the binding site on Cdc4 and a catalytic cysteine residue on Cdc34 within the SCF^Cdc4^ ubiquitin ligase dimer (Tang et al., 2007). These sites lie predominantly in non-helical regions (all except 53 and 84), which seems consistent with the prerequisite for Sic1 to simultaneously be docked on Cdc4 and reach the ubiquitination site on Cdc34.

### 2D inter-residue scaling maps

Scaling maps are two-dimensional representations of IDP structural propensities. Here, for each pair of residues in the Sic1 sequence, we calculated the average Cα – Cα distances in the optimized ensembles and normalized them to the respective distances in a random coil (RC) state (Figure 4). This type of analysis identifies regional biases for expansion (red) or compaction (blue). Alternatively, the inter-residue distances of the optimized ensemble can be normalized by the prior ensemble (see SI Fig. S5). Ensembles which agree well with the SAXS data (Fig. 4 A, D-F) have inter-residue distances that are overall more expanded than that of a RC, since 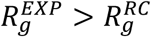 and 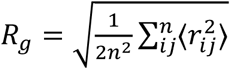. Conversely, the BME-optimized a99SBdisp and TraDES-SC ensembles (Fig. 4 B, C) are overall more compact. Compact regions near the diagonal indicate a propensity for secondary structure (see also Fig. 3). Since all ensembles agree with the FRET data between residues −1C and 90C, this region is similarly compact across all ensembles.

**Figure 4.**
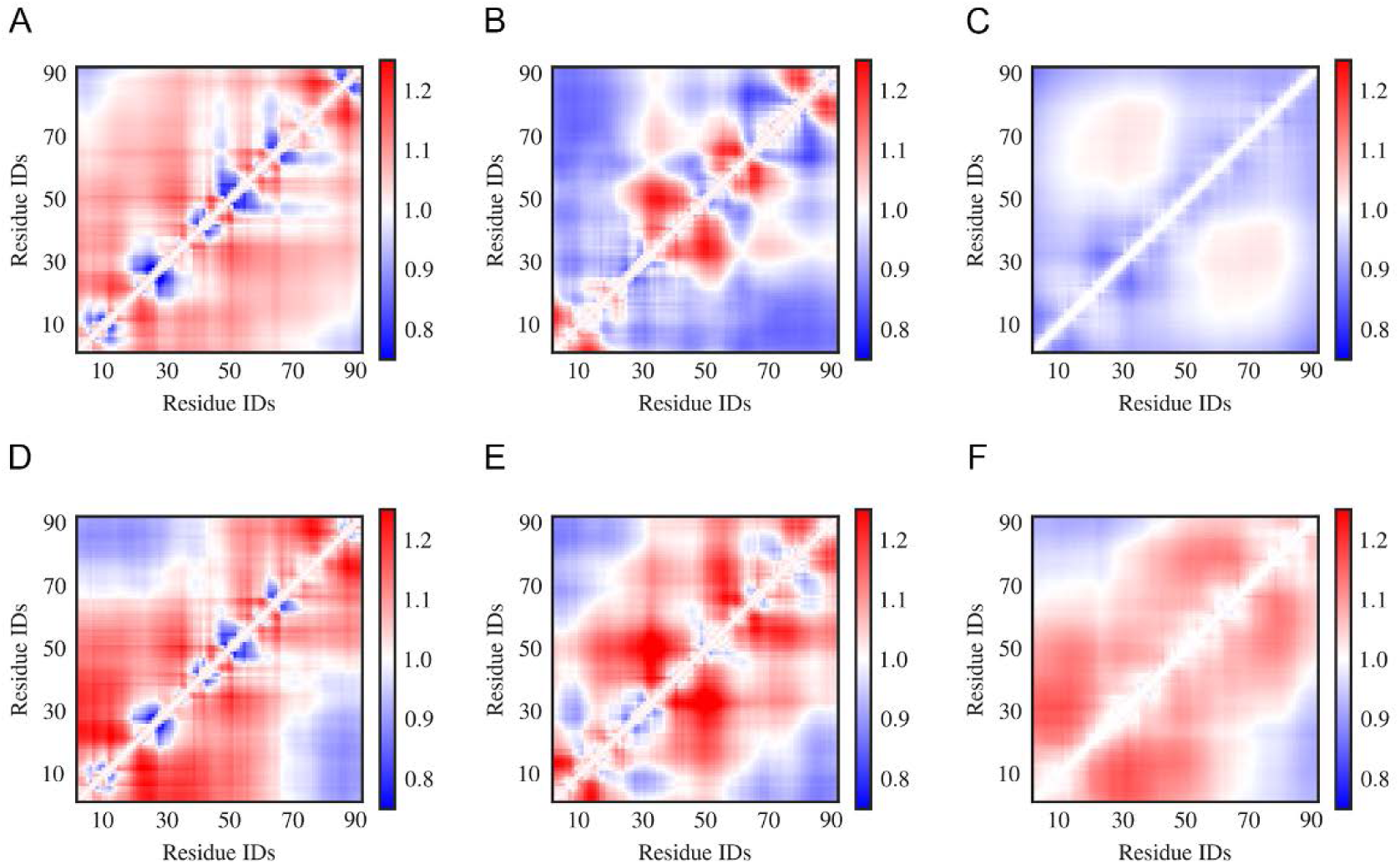
Inter-residue scaling maps of optimized Sic1 ensembles relative to a reference random-coil Sic1 ensemble (TraDES-RC). The ensembles were calculated using BME (**A-C**) or ENSEMBLE (**D-F**), and different initial ensembles: a03ws (**A, D**) a99SBdisp (**B, E**), and TraDES-SC (**C, F**). Ensemble-averaged distances between the Ca atoms of every unique pair of residues are normalized by the respective distances in a random coil. Regions in red are expanded relative to a random coil, while those in blue are more compact.

Notably, ENSEMBLE-optimization with different priors leads to different patterns of intermediate- and long-range distances, despite identical experimental restraints which included PRE measurements from six sites throughout the Sic1 sequence. This suggests, as Naudi-Fabra et al. have recently demonstrated (Naudi-Fabra et al., 2021), that multiple FRET and PRE measurements, which sufficiently sample the linear sequence of the protein, are needed to accurately reproduce intermediate- and long-range distances. Incorporating additional FRET restraints is expected to make ensembles optimized from different priors more similar in this respect. Indeed, examining the differences between the ensembles in Fig. 4 reveals inter-residue distances (i.e., possible FRET label locations) which will have the strongest discriminatory power between ensembles.

### Intra-chain interactions

Determining which specific molecular interactions determine the observed structural properties of IDP ensembles is an important goal. Knowledge of these interactions connect sequence properties to structural properties, allow testable predictions for the effects of mutations, and aid the rational design of molecules that bind disordered protein sequences with high affinity and specificity, stabilizing distinct IDP conformations (Ambadipudi and Zweckstetter, 2016; Robustelli et al., 2021). However, the experimental data, which are spatially and temporally averaged and are affected by noise, are insufficient to restrain distances and angles between groups of atoms, such that specific molecular interactions in conformations could be identified (e.g., hydrogen bond).

Including information from a force field which describes bonded and nonbonded interactions between the atoms, partially removes the degeneracy of the problem. The BME approach, which produces the minimum perturbation to the prior ensemble so that it fits the experimental data, is expected to retain the maximum amount of this information possible. Conversely, in the ENSEMBLE and similar approaches, which do not explicitly consider deviation from the prior ensemble, it is unclear in what capacity information about specific molecular interactions is retained. We therefore sought to compare the specific interactions in the resulting optimized ensembles. It is important to note that in our use of ENSEMBLE, PRE data was used as a restraint, whereas in our use of BME, PRE data was used as validation. This is expected to affect the inferred patterns of molecular interactions, in addition to the differences between optimization methods.

Excluding hydrophobic contacts, which are relatively non-specific, we hypothesized that the most likely interactions were hydrogen bonds (Fig. 5) and pi-contacts (Fig. 6). The Sic1 sequence has a high fraction of polar and charged residues (~54 %) that can participate in hydrogen bonding. Sic1 also has a high fraction of residues with sidechain pi bonds (~23%) and small residues with relatively exposed backbone peptide bonds (~52%) (Vernon et al., 2018). Prior to phosphorylation, Sic1 does not have any negatively charged residues, thus excluding salt-bridges and electrostatic attraction. To distinguish short-range/secondary structure (Fig. 3) from long-range tertiary contacts, we examine only those interactions with a sequence separation |*i* − *j*| > 10.

**Figure 5.**
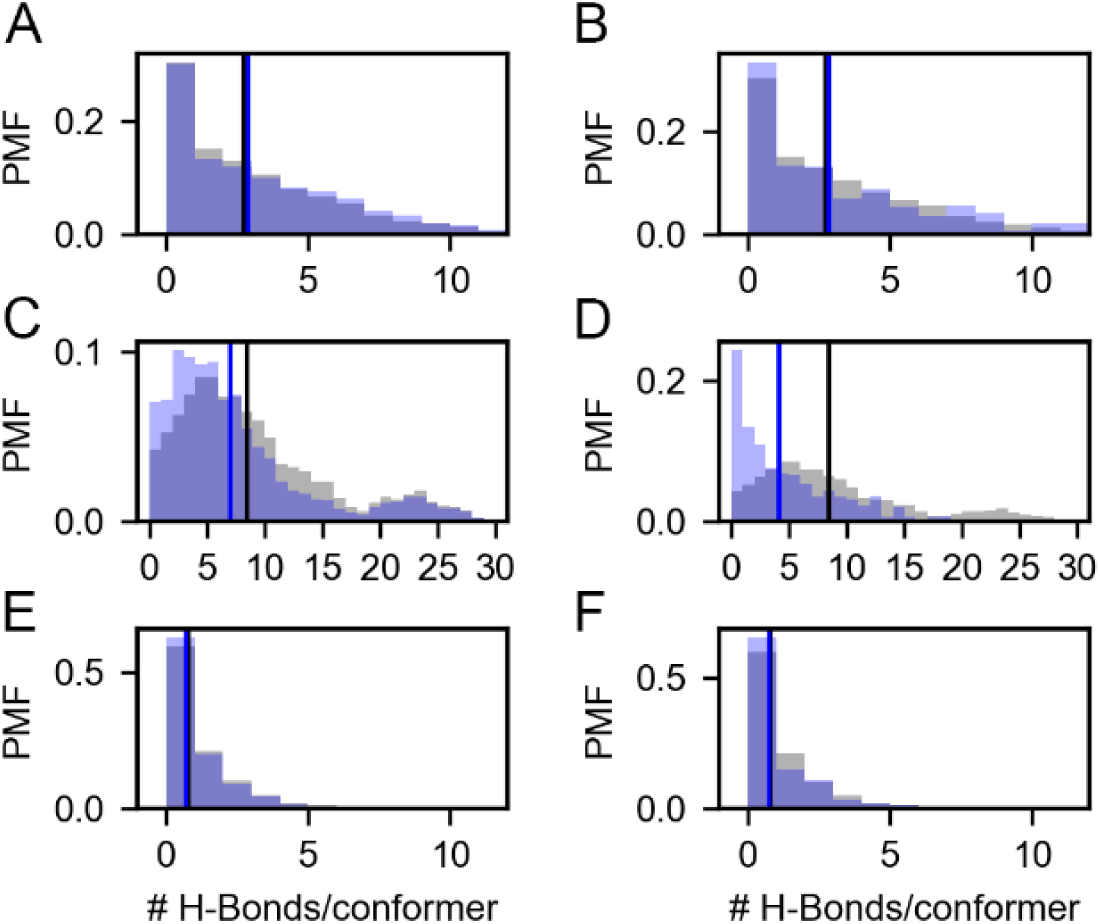
Probability mass functions (PMFs) of hydrogen bonds per conformer in prior (gray) and posterior (blue) Sic1 ensembles. The ensembles were calculated using BME (**A, C, E**) or ENSEMBLE (**B, D, F**), and different priors: a03ws (**A, B**) a99SBdisp (**C, D**), and TraDES-SC (**E, F**). Note that the x-axis for the a99SBdisp prior is larger than that of the a03ws and TraDES-SC priors.

**Figure 6.**
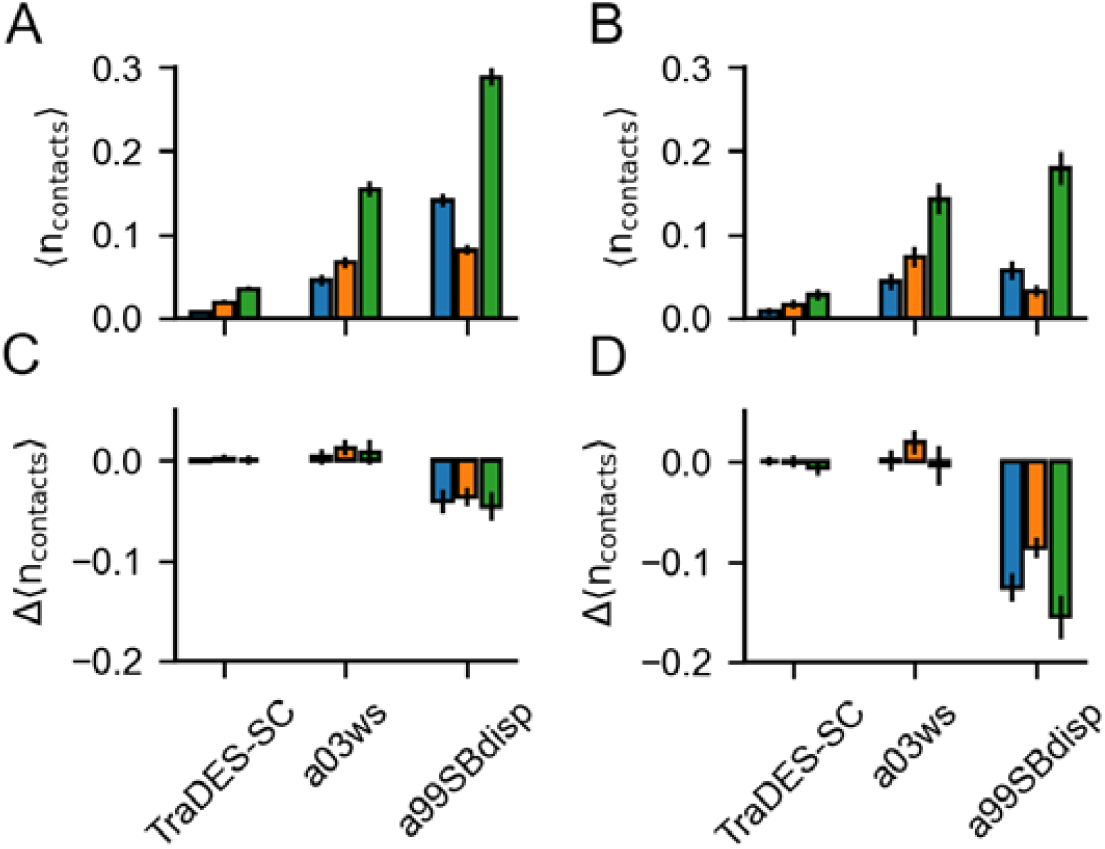
Pi-interactions in optimized Sic1 ensembles. Average number of pi-contacts per conformer using BME (**A**) and ENSEMBLE (**B**) and different priors. Pi contacts are separated into sidechain – sidechain (sc-sc, blue), back-bone – back-bone (bb-bb, orange) and sidechain – backbone (sc-bb, green). The change in the average number of pi-contacts per conformer induced by ensemble optimization via BME (**C**) and ENSEMBLE **(D**). Error bars are estimated by bootstrapping.

### Hydrogen bonds

Figure 5 shows the probability mass function (PMF) of the number of hydrogen bonds (H-bonds) per conformer in the prior (grey) and in the BME and ENSEMBLE optimized (blue) ensembles. H-bond contacts were defined using the distance and angle criteria established previously (Baker and Hubbard, 1984) and implemented in MDTraj (McGibbon et al., 2015). Vertical lines show the first moments of the corresponding PMFs. The TraDES-SC prior, for which there is no force-field describing non-bonded interactions, has very few H-bonds (i.e., ⟨*n*_*H*−*bonds*_⟩ = 0.7), corresponding to the small but finite probability of meeting the H-bond criteria by chance. As such, both BME and ENSEMBLE optimization have a very small effect on the H-bond propensity (Fig. 5 E, F).

As expected, there are significantly more H-bond contacts in the a03ws prior compared to the statistical noise in TraDES-SC (Fig. 5 A, B). Optimization using either BME or ENSEMBLE slightly increases the average number of H-bonds per conformer. This is consistent with the slight decrease in *R*_*g*_ and *R*_*ee*_, as the conformer *r*_*g*_ is inversely correlated with the number of H-bonds/conformer (see SI Fig. S6).

The a99SBdisp prior has an even higher average number of H-bonds/conformer than the a03ws prior, and the PMF is bimodal (Fig. 5 C, D). The differences in hydrogen bonding between a03ws and a99SBdisp may reflect parameterization choices in a99SBdisp to maintain accuracy for folded proteins (Robustelli et al., 2018). Alternatively, this may reflect incomplete sampling of extended structures, as simulations of Sic1 using enhanced-sampling techniques and the a99SBdisp force field produced *RR*_*gg*_ similar to that of a03ws and experiment (Shrestha et al., 2021). Interestingly, while a03ws has higher helical propensities than a99SBdisp and helical stability is largely driven by hydrogen bonding, it shows lower propensity for long-range H-bond interactions.

Whereas for a03ws both optimization methods result in qualitatively similar H-Bond PMFs, for a99SBdisp they differ considerably. Both optimizations reduce the average number of H-bonds per conformer; however, ENSEMBLE optimization removes the highly H-bonded subpopulation, and the resulting PMF is monotonically decreasing and similar to that of a03ws. Conversely, BME optimization retains this minor subpopulation, and shifts the center of the major subpopulation.

One reason for discrepant ENSEMBLE and BME H-bond inferences is how they achieve agreement with the SAXS data. The subpopulation of highly H-bonded conformations has a very compact radius of gyration (~2 nm, see SI Fig. S6) compared to the experimental radius of gyration (~ 3 nm). ENSEMBLE optimization prioritizes agreement with experimental data by eliminating the compact and highly H-bonded subpopulation. BME optimization seeks a balance between agreement with experiments and deviation from the prior, retaining this subpopulation at the expense of SAXS agreement, but smaller deviation from the a99SBdisp prior.

Experimental data is known to make ensembles more similar to one another (Ahmed et al., 2021; Larsen et al., 2020; Tiberti et al., 2015). Our results show that ensembles that agree with experimental data *and* were generated from an MD prior (a03ws-BME, a03ws-ENSEMBLE, a99SBdisp-ENSEMBLE) have similar H-bond PMFs. They are monotonically decreasing and have an average number of H-bonds per conformer ⟨*n*_*H*−*bonds*_⟩, between 3 and 4. However, our experimental data alone is insufficient to define H-bonds, as shown by the ENSEMBLE posterior ensembles (Fig. 5 B, D, F). Overall, this analysis suggests that the specific tertiary contacts and the nature of their molecular interactions in ensembles should be interpreted with caution. The type and amount of experimental data used here is insufficient, however incorporation of information about non-bonded interactions from MD force fields removes at least some of this degeneracy.

### Pi interactions

Although fixed charge atomistic MD force-fields do not explicitly include polarization and quantum effects to describe pi-interactions, they are valuable for understanding the relative importance of pi-interactions *vs*. other modes of interactions in stabilizing liquid-liquid phase separation in IDPs(Murthy et al., 2019; Schuster et al., 2020; Zheng et al., 2020). In folded protein structures, Vernon et al.(Vernon et al., 2018) found that the frequency of planar pi contacts strongly correlates with the quantity and quality of the experimental data and with the quality of the fit of the structure to the data. This suggests that current force fields may underestimate the relative importance of pi-pi interactions, and thus they appear more frequently when structures are more experimentally constrained. We therefore sought to determine: *(i)* whether the experimental data on Sic1 would refine the average number of planar pi-pi contacts per conformer in the ensembles and *(ii)* whether BME and ENSEMBLE optimization would result in different pi-pi contact frequencies.

Figure 6 shows the average number of planar pi-pi contacts per conformer, ⟨*n*_*pi*−*pi*_⟩,, in the optimized ensembles and the differences upon optimization. Pi contacts were defined using the distance and angle criteria presented by Vernon et al. (Vernon et al., 2018) and calculated using custom Python scripts provided by the authors. These contacts are classified as interactions between backbone amide groups (bb-bb), side chain amide, carboxyl, guanidinium or aromatic groups (sc-sc), or between backbone and side chain (bb-sc).

The TraDES-SC ensembles show the average number of each type of pi-contacts formed by chance, since there is no force field describing non-bonded interactions. Like for H-bonds, optimization of prior TraDES ensembles using either BME or ENSEMBLE did not change the frequency of pi-contacts. A03ws exhibits higher ⟨*n*_*pi*−*pi*_⟩ than TraDES-SC, suggesting that this force-field can somehow reproduce the pi-interaction geometries even without explicitly including polarization and quantum effects. Both BME and ENSEMBLE optimization did not significantly alter ⟨*n*_*pi*−*pi*_⟩. Like for H-bonds, the correlation between compactness, especially *r*_*g*_, with the number of contacts is the major driver for changes in ⟨*n*_*pi*−*pi*_⟩, upon optimization. In contrast to H-bonds, there are few conformations with more than one pi-interaction, and so the effect of optimization is more attenuated. Moreover, planar pi-pi interactions often involve groups with H-bond donors and acceptors, presenting an additional degree of degeneracy since the current experimental data do not directly report on pi-interactions.

Like for H-bonds, a99SBdisp has a higher average number of all types of pi-contacts than does a03ws and conformations with more than one pi-interaction are more compact (e.g., 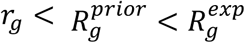). Consequently, both optimization methods significantly reduce the average number of pi-contacts. However, ENSEMBLE optimization, which results in better fits to the SAXS data, reduces ⟨*n*_*pi*−*pi*_⟩ more than BME optimization, which balances agreement with the SAXS data with deviation from the prior.

Intuitively, experimental data alone is insufficient to meaningfully describe pi-interactions in the absence of a force field (e.g., the TraDES-SC optimized ensembles). When the prior is constructed using a force field that describes the interaction geometries/strengths *and* the optimized ensembles agree with experimental data (e.g., BME-a03ws, ENSEMBLE-a03ws, and ENSEMBLE-a99SBdisp) the resulting ensembles have similar average numbers of pi-contacts (see also Fig. S7). As previously mentioned, experimental data makes the ensembles more similar, only when there exist interactions which can be re-weighted, and they are correlated with experimental data.

## SUMMARY AND CONCLUSIONS

Conformational ensembles for the disordered Sic1 protein were obtained by using experimental data (SAXS, CS, PRE and smFRET) as restraints and validation on three prior ensembles that were generated using either MD force fields or a statistical coil approach. The ensembles were optimized for agreement with the experiment using two contrasting modeling methodologies, Bayesian Maximum Entropy (Bottaro et al., 2020) (BME) and ENSEMBLE (Krzeminski et al., 2013). We compared the six different outcomes by examining global dimensions (e.g., *R*_*g*_), secondary structure propensities, inter-residue distances and specific non-local interactions, i.e., H-bonds and pi-interactions. Overall, differences between ensembles optimized using different priors were greater than when using the same prior with different optimization methods. Differences between methods were greatest when the priors were in poor agreement with experimental data, as BME balances perturbation of the prior ensemble with experimental agreement, whereas ENSEMBLE only focuses on the latter.

An advantage of MD priors is that they contain explicit information about specific molecular interactions (e.g., H-bonds and pi-interactions) that can be modulated, though not uniquely determined by experimental data. However, a disadvantage of MD priors is that they, by design and/or due to computational limitations, only sample a limited region of the entire conformational landscape. If incorrectly biased (e.g., overly compact) this will result in more significant re-weighting and experimental data may be insufficient to debias the ensemble. Future work would benefit from priors which are in better agreement with more than one type of experimental data prior to optimization.

Noting that ensembles optimized from different priors make different predictions regarding secondary structure, intermediate- and long-range distances, it appears that additional experimental data is needed, either as restraints or post-hoc validation. For secondary structure, this could include RDC data, which has been published for Sic1 but was not used in this analysis (Mittag et al., 2010, 2008), and fluorescence anisotropy decay, which reports on segmental dynamics of IDPs (Milles and Lemke, 2014). For intermediate- and long-range contacts, the *C*α – *C*α distance maps can be used to design maximally discriminating FRET label locations. Lastly, development of more rigorous ensemble optimization tools that integrate complementary biophysical data on multiple scales will lead to more accurate descriptions of conformational ensembles of IDPs and enable a mechanistic understanding of their biological function in implication in pathologies.

## METHODS

The SAXS and smFRET data from our group was recently published (Gomes et al., 2020) and the NMR data was published elsewhere (Mittag et al., 2010, 2008). The unoptimized MD ensembles (a03ws and a99SBdisp) were generated by resampling the original simulations (Robustelli et al., 2018) with a stride of 40 frames, resulting in Δ*t* = 7.2 ns between consecutive frames. For BME, forward calculation of the SAXS data was performed using Pepsi-SAXS (Grudinin et al., 2017) (see SI section 2); chemical shift data were calculated using ShiftX; smFRET data were calculated as described previously (Gomes et al., 2020). To accommodate smFRET measurements in BME, the J-Coupling module of BME was used, since both calculations involve a weighted linear average. The weight of the smFRET data relative to other data types was increased by scaling down the smFRET standard deviation by a factor of Ω^1/2^ (see SI section 1 for more details). For ENSEMBLE, calculations were performed as described previously (Gomes et al., 2020), using either the default conformer generation (TraDES-SC) or the resampled MD simulations (a03ws and a99SBdisp) as initial pools.

PRE intensity ratios were calculated using DEER-PREdict (Tesei et al., 2021) v0.1.8 with an effective correlation time of the spin label of *τ*_*C*_ = 2 ns, *τ*_*t*_ = 0.5 ns, total INEPT time *t*_*d*_ = 10 ms, reduced transverse relaxation rate *R*_2_ = 10 Hz, and proton Larmor frequency *ω*_*H*_/2*π* = 500 MHz. The root-mean-squared error between the calculated and experimental intensity ratios was calculated for each label location (−1, 21, 38, 64, 83, and 90) and the final PRE validation score is the root-mean-squared average of the six RMSDs.

*Analysis* of optimized and unoptimized ensembles (radius of gyration, scaling maps, DSSP, H-Bonds) were performed using MDTraj (McGibbon et al., 2015) v1.9.5. Pi-contact analysis was performed using scripts provided by Vernon et al. (Vernon et al., 2018) Uncertainties in the secondary structure propensities, and in the average number of each type of pi-contact were determined using bootstrapping; i.e., the calculations were performed on *N* conformations randomly sampled (with replacement) from the initial ensemble, with either uniform weights (*w*_*i*_ = 1/*N*) for calculations on the prior ensembles or the ENSEMBLE-optimized ensembles, or with the BME determined weights 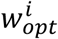 for the BME-optimized ensemble. The uncertainties were calculated as the standard deviation of the parameter of interest for the *B* = 1000 bootstrapped ensembles.

## ASSOCIATED CONTENT

### Supporting Information

Optimal weighting of smFRET data in BME, calculation of SAXS data and hydration effects, radius of gyration distribution before/after optimization in BME, 2D contact maps scaled by the prior, 2D histograms of radius of gyration vs. number of hydrogen bonds, differences in pi contacts between optimized ensembles.

## AUTHOR INFORMATION

### Author Contributions

G.-N.G. and C.C.G. designed and coordinated the research, and wrote the manuscript. A.N. and G.-N.G. performed data analysis and made figures and tables for the manuscript. All authors have given approval to the final version of the manuscript.

### Funding Resources

This work has been supported by the Natural Sciences and Engineering Research Council of Canada (NSERC RGPIN 2017 – 06030 to C.C.G.).

## ACKNOWLEDGMENT

The authors are grateful to Dr. J.-D. Forman-Kay from Sick Kids Hospital for providing the NMR data of Sic1 used in this study and to Dr. M. Krezminski from her lab for his support in using the ENSEMBLE program. We thank Dr. T. Mittag and Dr. E. Martin from St. Jude Hospital for providing SAXS data of Sic1. The authors are grateful to Dr. D.E. Shaw for providing molecular dynamics simulation data of Sic1. We also thank Dr. K. Lindorff-Larsen and Dr. S. Bottaro for technical help with using the BME program.

