## Supplemental Material for "Integrative conformational ensembles of Sic1 using different initial pools and optimization methods"

**Contents:** Optimal weighting of smFRET data in BME, calculation of SAXS data and hydration effects, radius of gyration distribution before/after optimization in BME, 2D contact maps scaled by the prior, 2D histograms of radius of gyration vs. number of hydrogen bonds, differences in pi contacts between optimized ensembles

### 1 Relative weight of smFRET vs. SAXS and CS in BME optimization – finding Ω

The information content is SAXS is greatly overestimated if $\chi^2_{SAXS}$ is weighted by the number of SAXS data points ($m_{SAXS} \approx 200$). The information content of SAXS data is usually estimated on the basis of Shannon's sampling theorem[1] (i.e., $N_s$, the number of Shannon channels) or other information theoretic approaches[2] (e.g., $N_g$, the number of 'good parameters' as defined in Ref[2]). For the Sic1 experimental SAXS data, $N_s = 7.3$ and $N_g = 4$ were determined using BayesApp[3] suggesting that the SAXS data is dramatically oversampled. Moreover, SAXS data report purely statistical errors, whereas systematic errors, for example in background subtraction are typically unknown. Similarly, commonly used implicit solvent SAXS forward calculators require specification of free parameters describing the contribution of the hydration layer and displaced solvent[4,5] introducing additional uncertainty.

Taken together, this suggests that the relative weight of SAXS data to smFRET data is overestimated if it is taken as $m_{SAXS}/m_{FRET} \approx 200$, where $m_{FRET} = 1$ (smFRET measurement on one pair of residues). Similar arguments can be made for the CS data, wherein calculation errors and statistical independence of the data are difficult to estimate[6], making the absolute value of $\chi^2_{CS}$ relative to other data types of limited use.

Analogous to $\theta$, we introduce a hyperparameter Ω such that $\chi^2_{TOTAL} = \chi^2_{SAXS} + \chi^2_{CS} + \Omega\chi^2_{FRET}$. In theory, the relative weights of all three experiments could be varied, though for simplicity we only vary the weight of smFRET relative to SAXS and CS data. This is justified since $m_{SAXS} \approx m_{CS} \approx 200$ data points and since SAXS and CS report on properties that are only weakly correlated[7–9].

To determine Ω, we perform BME optimization (including $\theta$ tuning) for fixed values of Ω from 1 to 400 and examine how the choice of Ω affects $\chi^2_{SAXS}$, $\chi^2_{FRET}$, the PRE validation score and the amount of reweighting ($N_{eff}$) at the chosen optimum $\theta$. For a03ws the optimum $\theta$ is determined from the minimum of the PRE validation score. For a99SBdisp and TraDES-SC (data not shown), for which the PRE validation score monotonically worsens with increased reweighting, we use the knee of the $\chi^2_{TOTAL}$ graph to determine the optimum $\theta$. For all ensembles, examining $\chi^2_{SAXS}$, $\chi^2_{FRET}$, the PRE validation score, and $N_{eff}$ at the optimum $\theta$ shows that increasing Ω from 1 to $\Omega \approx 75$ improves the fit to the smFRET data and the PRE validation data, without negligible deterioration of the SAXS fit, and with only marginally more reweighting.

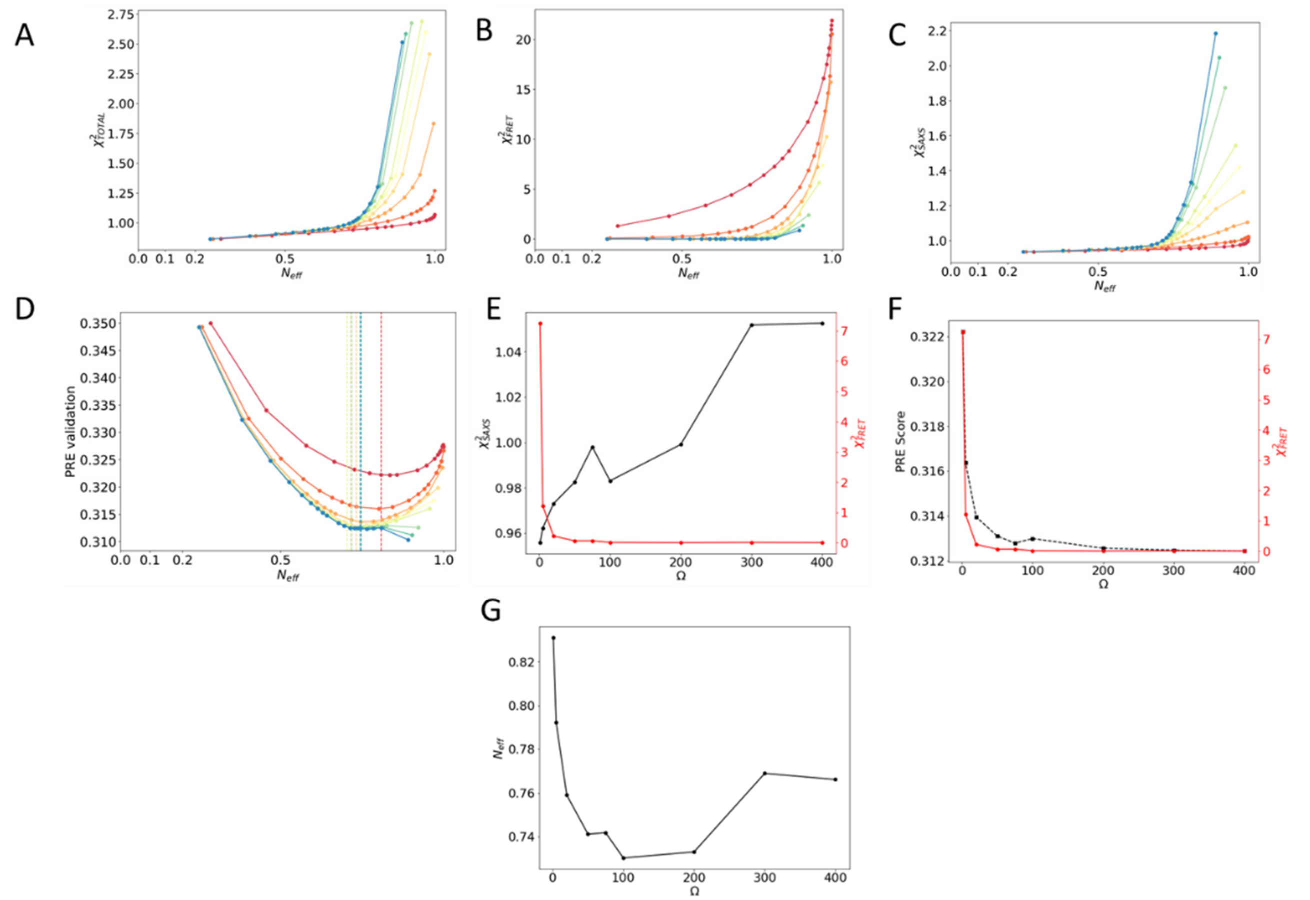


**Supplementary Figure 1:** Determining the hyperparameter $\boldsymbol{\Omega}$ for the a03ws BME optimization. (**A-C**) The hyperparameter $\boldsymbol{\Omega}$ was varied from $\boldsymbol{\Omega = 1}$ (red) to $\boldsymbol{\Omega = 400}$ (blue) and for each fixed value of $\boldsymbol{\Omega}$ the hyperparameter $\boldsymbol{\Theta}$ was varied (resulting in lower $\boldsymbol{N_{eff}}$ at lower $\boldsymbol{\theta}$, greater reweighting to agree with experiment). (**D**) For each fixed $\boldsymbol{\Omega}$, $\boldsymbol{\theta}$ was determined from the minimum of the PRE validation. (**E-F**) At the above determined optimum $\boldsymbol{\theta}$ the behavior of $\boldsymbol{\chi^2_{SAXS}}$ (**E**) or PRE validation (**F**) and $\boldsymbol{\chi^2_{FRET}}$ as a function of $\boldsymbol{\Omega}$. At $\boldsymbol{\Omega \approx 75}$ there is substantial improvement in $\boldsymbol{\chi^2_{FRET}}$ with negligible deterioration of $\boldsymbol{\chi^2_{SAXS}}$. At $\boldsymbol{\Omega \approx 75}$ both the PRE validation score and $\boldsymbol{\chi^2_{FRET}}$ improve relative to $\boldsymbol{\Omega = 1}$. (**G**) As $\boldsymbol{\Omega}$ increases $\boldsymbol{N_{eff}}$ decreases, however $\boldsymbol{\Omega \approx 75}$ represents only marginally more reweighting (lower $\boldsymbol{N_{eff}}$) compared to $\boldsymbol{\Omega = 1}$.

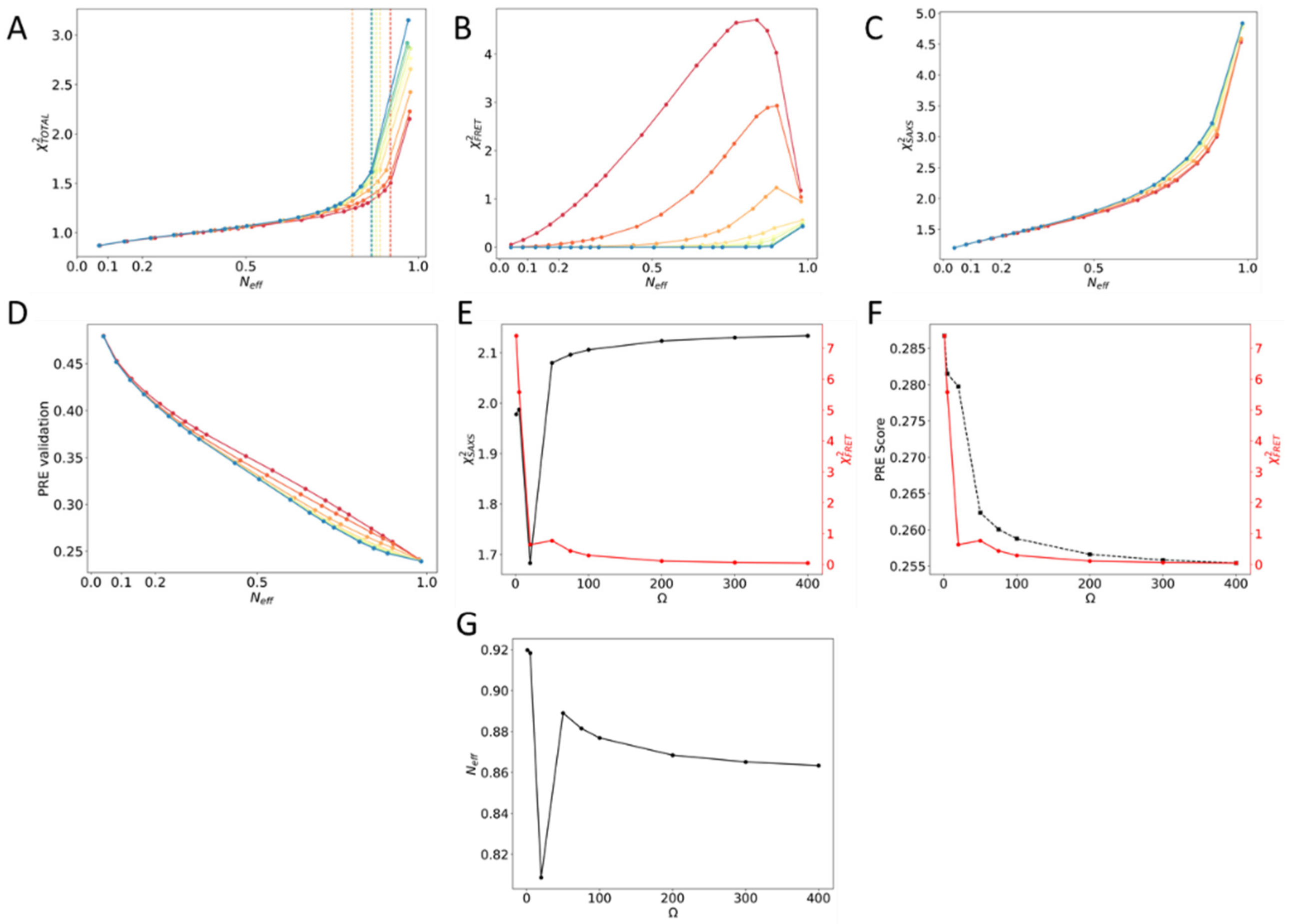


**Supplementary Figure 2:** Determining the hyperparameter $\boldsymbol{\Omega}$ for the a99SBdisp BME optimization. (**A-C**) The hyperparameter $\boldsymbol{\Omega}$ was varied from $\boldsymbol{\Omega = 1}$ (red) to $\boldsymbol{\Omega = 400}$ (blue) and for each fixed value of $\boldsymbol{\Omega}$ the hyperparameter $\boldsymbol{\Theta}$ was varied (resulting in lower $\boldsymbol{N_{eff}}$ at lower $\boldsymbol{\theta}$, greater reweighting to agree with experiment). (**A**) For each fixed $\boldsymbol{\Omega}$, $\boldsymbol{\theta}$ was determined from the knee of the $\boldsymbol{\chi^2_{TOTAL}}$ graph. **(E-F)** At the above determined optimum $\boldsymbol{\theta}$ the behavior of $\boldsymbol{\chi^2_{SAXS}}$ (**E**) or PRE validation (**F**) and $\boldsymbol{\chi^2_{FRET}}$ as a function of $\boldsymbol{\Omega}$. At $\boldsymbol{\Omega \approx 75}$ there is substantial improvement in $\boldsymbol{\chi^2_{FRET}}$ with negligible deterioration of $\boldsymbol{\chi^2_{SAXS}}$. At $\boldsymbol{\Omega \approx 75}$ both the PRE validation score and $\boldsymbol{\chi^2_{FRET}}$ improve relative to $\boldsymbol{\Omega = 1}$. (**G**) As $\boldsymbol{\Omega}$ increases $\boldsymbol{N_{eff}}$ decreases, however $\boldsymbol{\Omega \approx 75}$ represents only marginally more reweighting (lower $\boldsymbol{N_{eff}}$) compared to $\boldsymbol{\Omega = 1}$.

### 2 SAXS forward calculation and implicit hydration effects

In ENSEMBLE, SAXS curves are calculated from conformers using CRYSOL[10] with the default hydration parameters. In BME, we use Pepsi-SAXS[11] with the default effective atomic radius $r_0$ and with the contrast of the hydration layer $\delta\rho$ determined as described in Figure captions. A value of $\delta\rho =$ 20 e/nm$^3$ is nearly optimum for all three prior ensembles upon reweighting. In principle, both $r_0$ and $\delta\rho$ could be further optimized using the iterative procedure outlined by Pesce and Lindorff-Larsen[12].

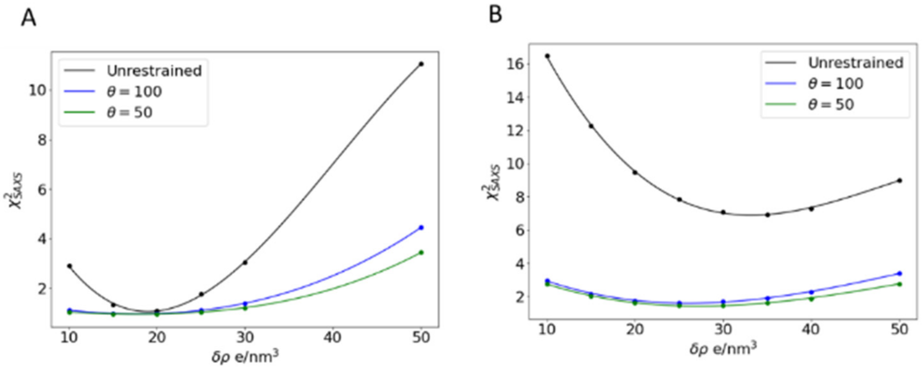


**Supplementary Figure 3:** Dependence of unoptimized and optimized ($\boldsymbol{\theta = 100}$ ***or*** $\boldsymbol{\theta = 50}$) $\boldsymbol{\chi^2_{SAXS}}$ on the hydration layer contrast $\boldsymbol{\delta\rho}$. A value of $\boldsymbol{\delta\rho}$=20 e/nm$^3$ is near the minimum for both a03ws and a99SBdisp optimized ensembles.

### 3 Distributions of radius of gyration for prior and BME-optimized ensembles

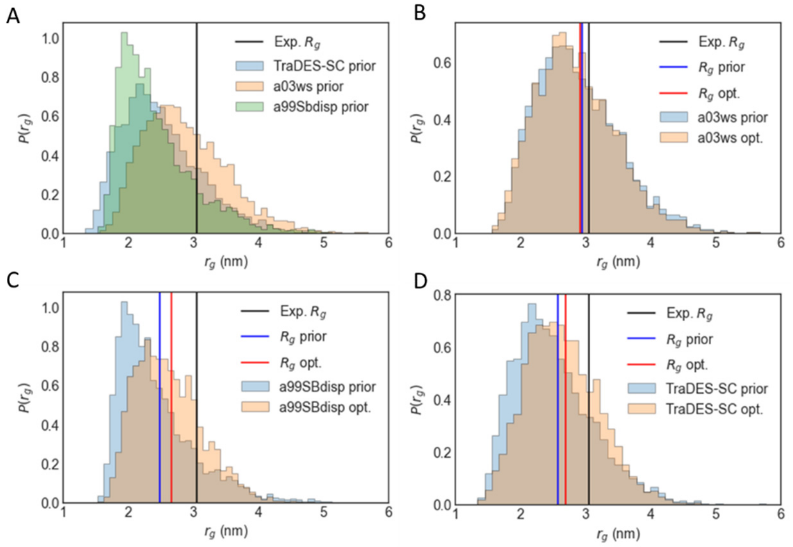


**Supplementary Figure 4:** Radius of gyration distributions before and after BME reweighting for the three prior ensembles. The experimental radius of gyration from Gomes et al., [9] is shown in black. The ensemble root mean squared radius of gyration is calculated as $\boldsymbol{R_g = \langle r_g^2 \rangle^{1/2}}$ either without (prior) or with (optimized) the BME determined weights. Note that the experimental $\boldsymbol{R_g}$ contains an additional contribution from solvent scattering, whereas the ensemble $\boldsymbol{R_g}$ is calculated from the protein coordinates using MDTraj[13] (see also Ref [12]).

### 4 Contact maps scaled by the prior

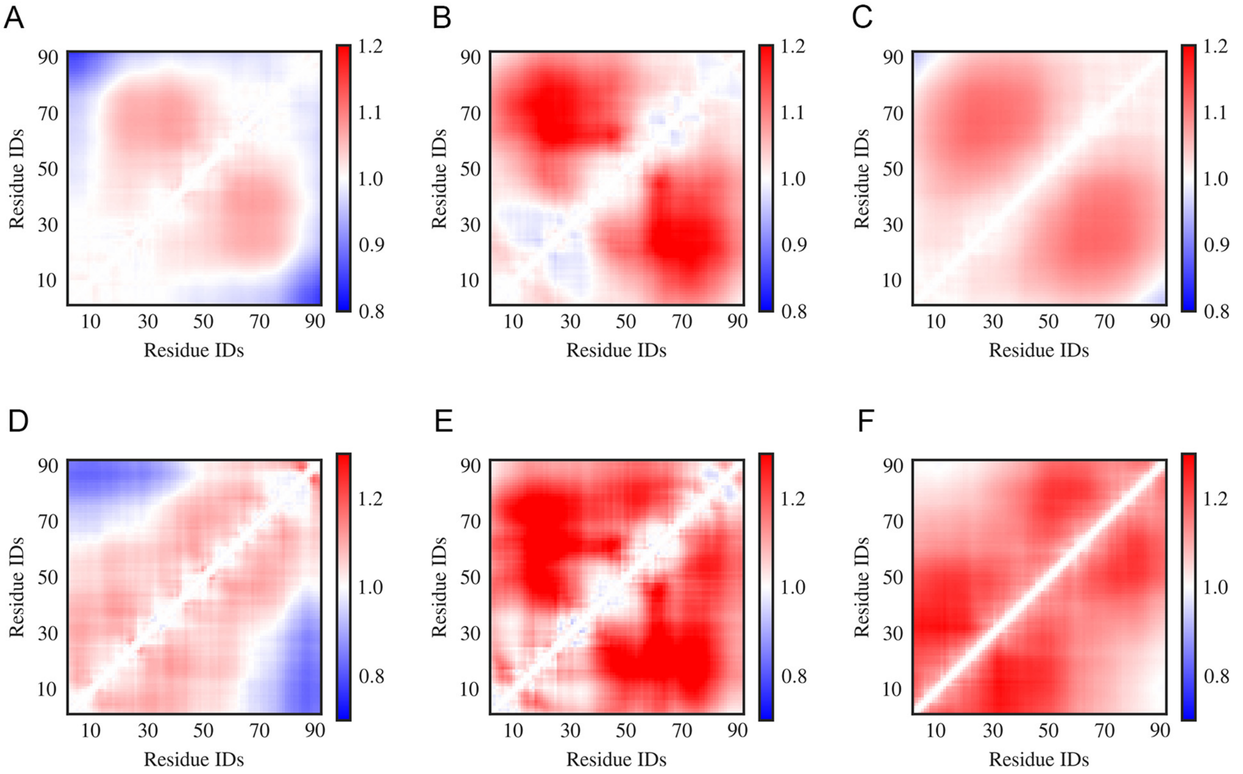


**Supplementary Figure 5:** Inter-residue scaling maps of optimized Sic1 ensembles relative to their respective prior ensembles. The ensembles were calculated using BME (**A-C**) or ENSEMBLE (**D-F**), and different initial ensembles: *a03ws* (**A, D**) *a99SBdisp* (**B, E**), and *TraDES-SC* (**C, F**). Ensemble-averaged distances between the $C\alpha$ atoms of every unique pair of residues are normalized by the respective distances in the prior ensembles. Regions in red are expanded relative to a random coil, while those in blue are more compact.

### 5 Radius of gyration vs. H-bonds

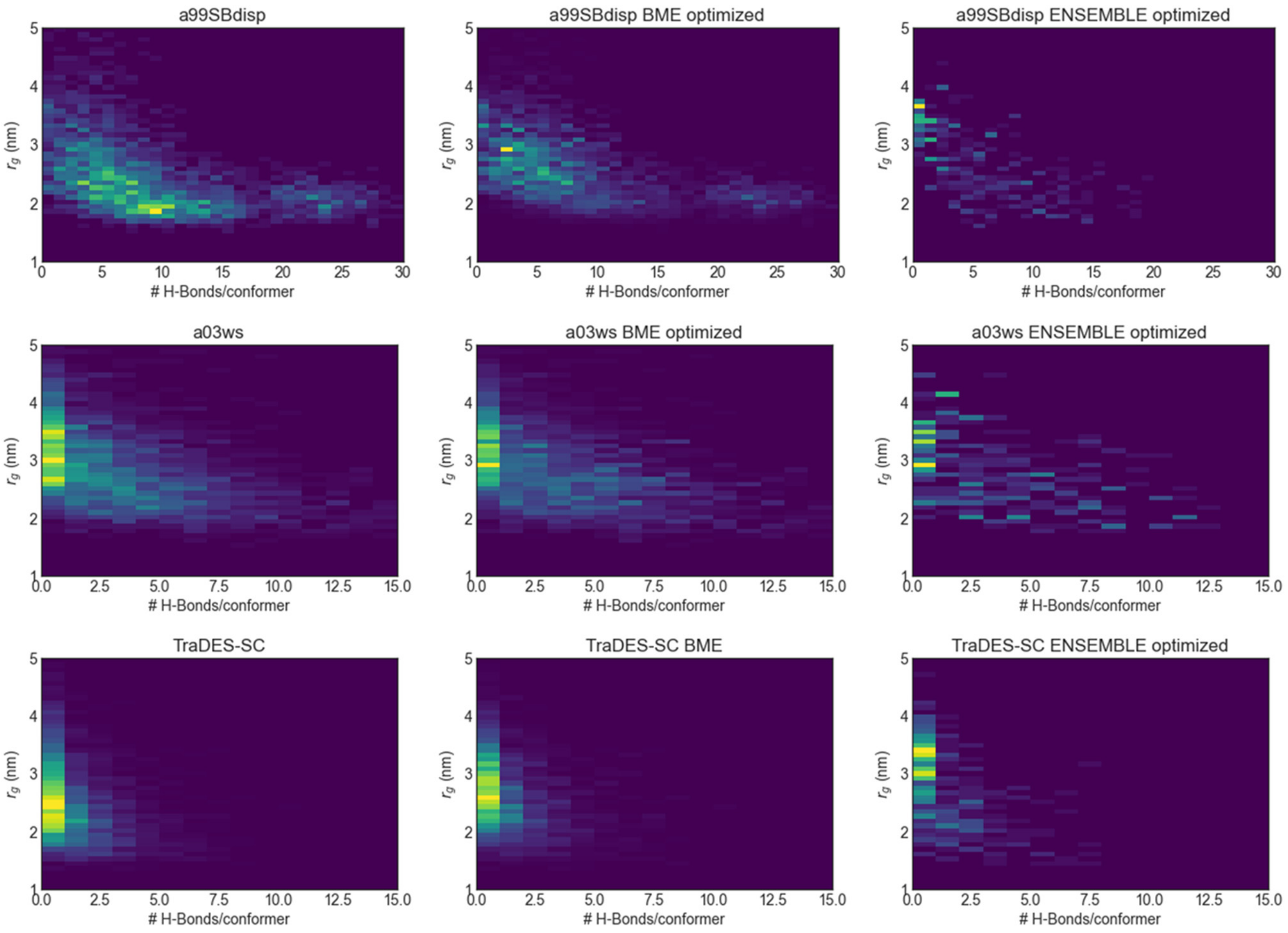


**Supplementary Figure 6**: 2D histograms showing the correlation between the radius of gyration vs. # of H-bonds/conformer for initial and optimized ensembles, using BME and ENSEMBLE on three different priors. Generally, more highly hydrogen bonded conformations have more compact radii of gyration.

### 6 Differences in pi contacts between optimized ensembles

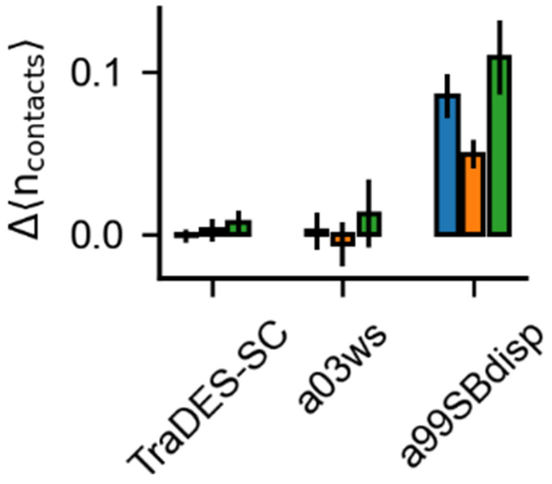


**Supplementary Figure 6:** Difference in number of Pi-interactions between Sic1 ensembles optimized by BME or ENSEMBLE. Average number of pi-contacts per conformer using BME subtracted by respective quantity using ENSEMBLE. Pi contacts are separated into *sidechain – sidechain* (sc-sc, blue), *back-bone – back-bone* (bb-bb, orange) and *sidechain – backbone* (sc-bb, green). Error bars are estimated by bootstrapping.
